# Melanopsin-mediated signals in natural and human-made environments

**DOI:** 10.1101/2025.06.07.658429

**Authors:** Pablo A. Barrionuevo, Francisco Diaz Barrancas

## Abstract

Melanopsin-expressing intrinsically photosensitive retinal ganglion cells (ipRGCs) play a critical role in regulating physiological and behavioral responses to light. However, little is known about how melanopsin and ipRGC signals are shaped by the statistical properties of real-world environments. Here, we analyzed statistics of melanopsin, ipRGC codification of extrinsic and intrinsic photoresponses, and luminance using hyperspectral images of natural and human-made scenes under daylight illumination. The statistics were obtained simulating receptive fields from current knowledge about ipRGCs anatomy and physiology. Our findings reveal that human-made environments exhibit significantly higher melanopsin, luminance, and ipRGC excitations compared to natural environments. In natural scenes, luminance contrasts were higher than melanopsin and ipRGC contrasts across most of the range. Melanopsin contrast was largely independent of excitation, and was significantly reduced for larger receptive fields. Differences between ipRGC codification models suggest an interaction between input weighting and environmental structure. These results indicate that modifications of natural regularities by human-made environments could affect ipRGC-driven physiology in everyday life.

## Introduction

The natural environment contains regularities that are crucial for the evolution of sensory physiological mechanisms. Through natural selection, an organism’s perceptual systems are strongly linked to the properties of its physical environment^1^. In humans, this connection is achieved through a combination of adaptive changes throughout life and inherent adaptations present at birth. These lifetime adaptive changes adjust the perceptual systems, for example, to increase sensitivity to regularities^2^. In addition to lifetime constraints, the design of perceptual systems is influenced by the specific tasks that organisms evolved to efficiently represent the variability of the natural world. Natural image statistics are a property of environments and they can be used to predict how neural responses should vary to efficiently encode them^3^. For example, the combination of excitatory and inhibitory signals in postreceptoral pathways are tuned to the principal components that explain most of the variability in natural images^4,5^. Especially important for sensory functions are the mean light intensity in the environment and the contrast, which is related to the spatial and temporal differences of light inputs^6^. These statistics have motivated different theories about visual perception, for example, regarding lightness perception^7^. However, most of these studies only dealt with photopic luminance, which is by definition driven by cones and weighted by the photopic sensitivity function “V (λ)” (Fig. 1A, inset). For example, Frazor and Geisler have analyzed luminance and luminance contrast driven by natural images during typical saccadic inspection^8^. Balboa and Grzywacz have shown that differences in the distributions of luminance contrast driven by different habitats are associated with receptive field sizes^9^. However, how the conclusions based on cone-driven responses can be extrapolated to other photoresponses is still unknown, particularly when receptive field have different sizes and spectral sensitivities are shifted.

**Figure 1.**
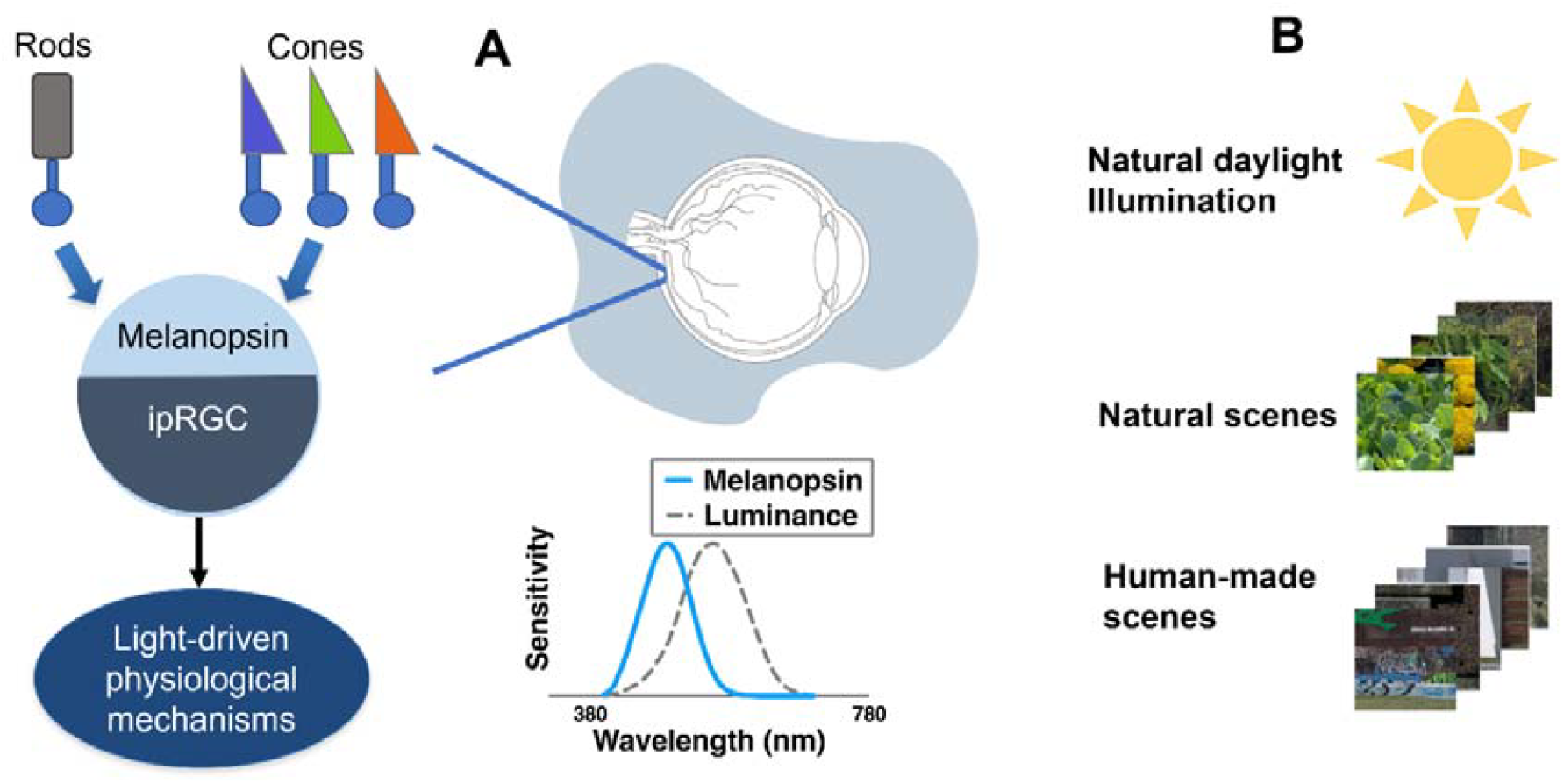
IpRGCs in the physical environments. A) Schematic representation of ipRGCs inferences and efferences. The photoreceptors are located in the retina of the eye reached by the lighting environment. The melanopsin sensitivity is spectrally shifted from the luminance (L+M cones) sensitivity (inset). B) Hyperspectral images were divided between natural and human-made environments obtained under natural daylight illumination. Images were obtained from a publicly available repository^48^ and represent the scenes used in this study (Table S1).

At the beginning of this century, it was demonstrated that a group of retinal ganglion cells were intrinsically photosensitive (ipRGCs) in mice^10,11^, and then in primates including humans^12,13^. These cells express the photopigment melanopsin^14^ (Fig. 1A). The light intensity range where melanopsin is active roughly overlaps with that of cones^12^. The role of these cells in the photoentrainment of the circadian rhythm was early discovered ^10,15^. Based on accumulated evidence, now, there is a consensus about the role of ipRGCs to provide light exposure information to several physiological functions for regulatory purposes and synchronization with the light environment^16^. These functions are pupillary control^17–19^, photophobia^20^, pain modulation^21^, melatonin secretion^22^, development^23^, activity^24^, mood^25^, alertness^26^, visual perception^27–31^, and cognition^32^. Since, ipRGCs are highly specialized to codify irradiance in primates^33^, and their impact in physiology is substantial, it is important to understand how these cells process the regularities in different visual environments.

IpRGCs are anatomically different than the parasol ganglion cells which are involved in the luminance codification^34^. The size of the ipRGCs dendritic field increases with eccentricity^12,35^. In comparison with ganglion cells involved in visual processes, ipRGCs have a larger dendritic field than parasol and midget ganglion cells throughout the retina^12^. As other ganglion cells^36^, IpRGCs are denser in the parafoveal region and its density decrease with periphery with a scotoma in the foveal pit^35,37^. There exist at least two populations of ipRGCs in humans (outer and inner), and the dendritic fields of these populations overlap^35^. In the human retina, parasol cells are 10% of the total ganglion cells^38^, instead ipRGCs percentage is around 0.4%^39^. Outer ipRGCs in humans provide a unique codification of cone signals, being L- and M-cone ON and S-cone OFF^12^. This S-cone OFF input is mediated by an S-cone amacrine cell that receives excitatory inputs from bipolar cells and provides inhibitory signals to ipRGCs^40^. This unique opponency has been confirmed in human pupillary measurements ^41–43^. While, parasol cells codify cone signals additively^44^. Another interesting characteristic of outer ipRGCs is that they don’t have spatial opponency^12^. Spatial opponency is fundamental in other visual ganglion cells to allow codification of borders^45^. However, large spatial contrasts could be detected by ipRGCs when comparing the outputs of adjacent cells; as it was demonstrated recently to obtain the melanopsin contrast sensitivity function^28^.

There is a close match between natural statistics and processing of contrasts in the early visual system, however most of the studies were carried out considering luminance (cone) statistics which are useful for visual purposes but other physiological functions are more affected by ipRGCs than parasol (luminance) ganglion cells. This work aimed to study the excitation (absolute values) and contrast statistical regularities of melanopsin and ipRGC codification in natural and human-made scenes.

## Results

We used hyperspectral scenes obtained in Portugal in previous studies ^46,47^ (Fig. 1B). Our analyses were based on anatomical and physiological features of human ipRGCs and parasol cells (see Materials and Methods section).

### Absolute values

We first assessed the absolute values of melanopsin, ipRGC codification (L M Mel ON/S OFF) and luminance (Fig. 2). For the three metrics, the human-made environments generated significantly higher absolute values than natural environments (Melanopsin: *t* = −5.4, *p* < 0.0001; Luminance: *t* = −3.67, p < 0.001; ipRGC: *t* = −4.3, *p* < 0.0001). The distribution spread didn’t differ from each other, since there were no differences between variances of the two data groups (Tables 1, S1). Median and interquartile values are summarized in Table 1.

**Table 1.** Summary of the median and interquartile values of the first-order (absolute) and second-order (contrast) statistics. Statistical differences between the two samples’ mean and variance from the transformed variable are also indicated.

|  | Melanopsin |  |  |  | ipRGC |  |  |  | Luminance |  |  |  |
| --- | --- | --- | --- | --- | --- | --- | --- | --- | --- | --- | --- | --- |
|  | Absolute |  | Contrast |  | Absolute |  | Contrast |  | Absolute |  | Contrast |  |
|  | Median | IQR | Median | IQR | Median | IQR | Median | IQR | Median | IQR | Median | IQR |
| Natural | 0.43 | 1.27 | 0.37 | 0.43 | 1.22 | 3.16 | 0.37 | 0.42 | 497 | 1547 | 0.46 | 0.47 |
| Human-made | 0.99 | 1.89 | 0.38 | 0.47 | 2.14 | 4.02 | 0.39 | 0.48 | 851 | 2109 | 0.44 | 0.54 |
| Differents? | Yes*** | No | No | No | Yes*** | No | No | No | Yes*** | No | No | No |

\*\*\* Differences between groups indicated for $p < 0.001$ . Statistics values are shown in Table S1.

**Figure 2.**
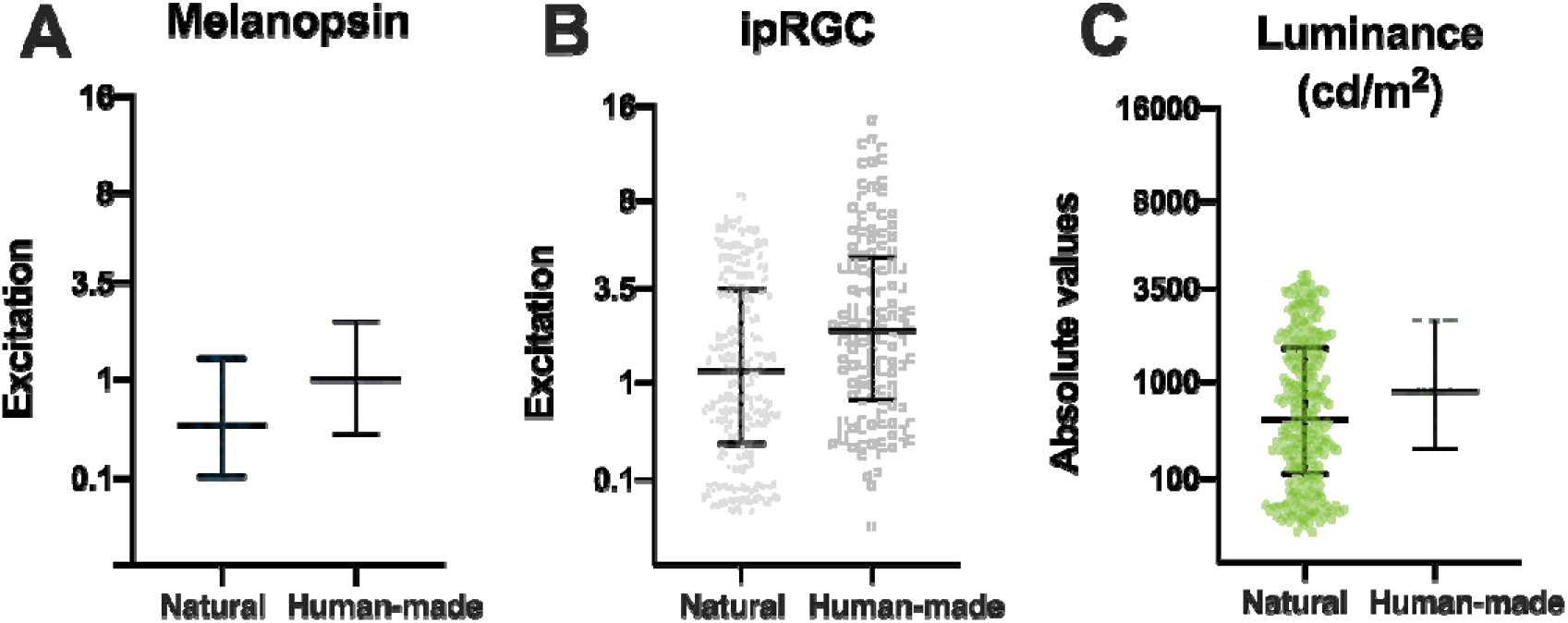
Absolute excitation values across visual channels and environments. (A) Distribution of melanopsin excitation values for natural and human-made environments. (B) ipRGC 1 excitation values based on weighted cone and melanopsin inputs. (C) Luminance values computed using the CIE 10° V(λ) function. In all three cases, human-made environments exhibit significantly higher absolute excitation values than natural environments (p < 0.001). Wide middle horizontal line represents median and error bars represent the interquartile range.

### Contrasts

We computed the absolute contrast between the simulated receptive fields in each scene (see Methos section). For the three metrics, no significant differences were found between human-made scenes and natural scenes for mean and variance values (Tables 1 and S2). Since the contrast is a relative variable, we could compare between different metrics. This contrast computation revealed that for natural environments (Fig. 3A), luminance contrasts are higher than melanopsin and ipRGCs signals [*F*(2, 949) = 4.78, *p* < 0.01; Tukey-Kramer post-tests: *p* = 0.99 (Melanopsin vs ipRGC), *p* < 0.05 (Melanopsin vs Luminance), *p* < 0.05 (Luminance vs ipRGC)]. However, these differences were not found for human-made environments (Fig. 3B; *F*(2, 459) = 1.28, *p* = 0.28). When contrasts of luminance and melanopsin are compared considering each receptive field (Fig. 3C and D), we found a differential behavior regarding the contrast magnitude. Luminance contrasts are higher than melanopsin contrasts only in the upper range, however for the low range this difference disappeared. This finding is similar for both natural scenes (Fig. 3C, luminance > melanopsin for contrasts higher than 0.25) and human-made scenes (Fig. 3D, luminance > melanopsin for contrasts higher than 0.29). We have also found that the overlapping between contrast clusters of these metrics is high (natural: *r* = 0.68, *p* < 0.0001; human-made: *r* = 0.77, *p* < 0.0001).

**Figure 3.**
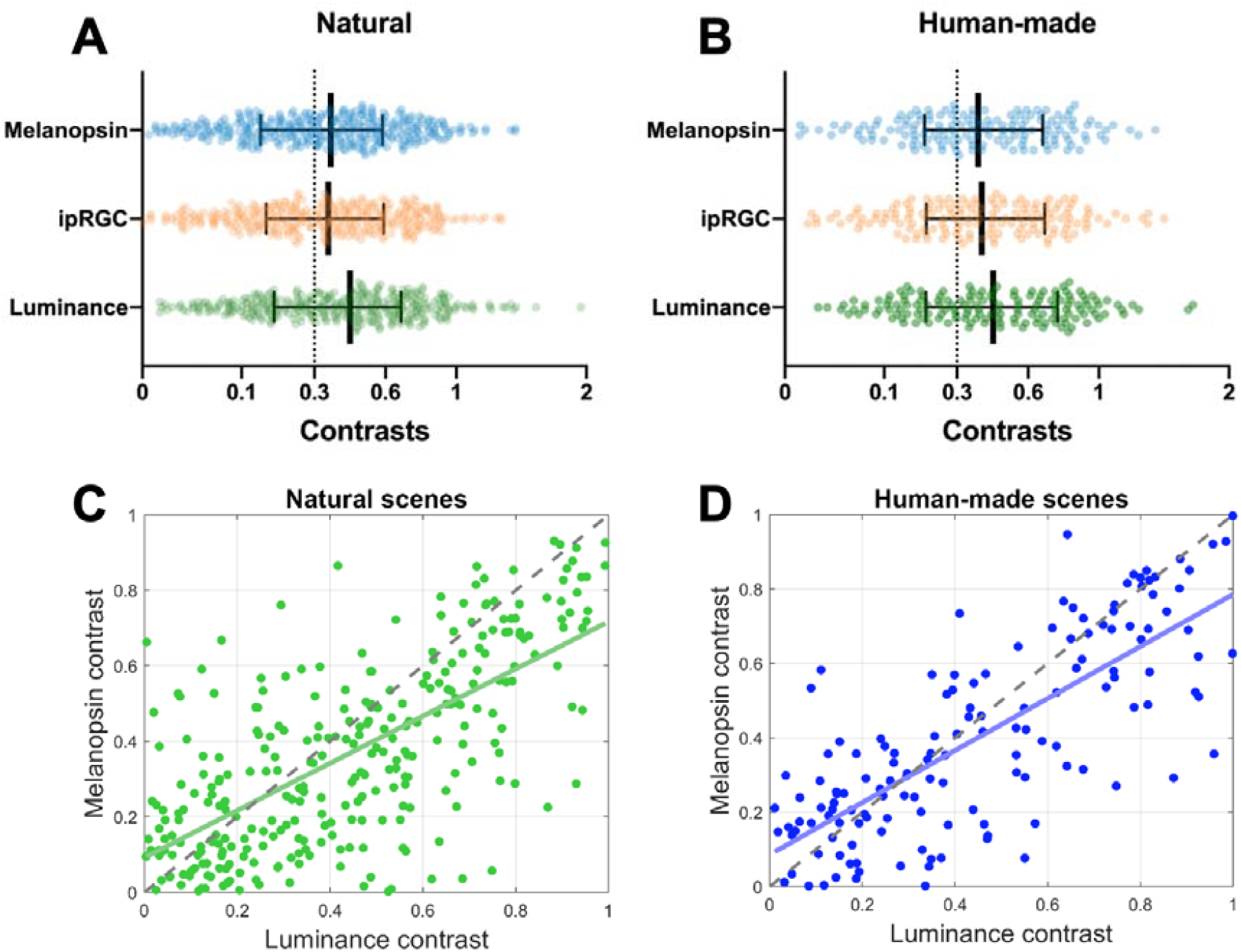
Contrast distribution in natural and human-made environments. (A) Contrast values for melanopsin, ipRGC codification, and luminance in natural scenes. (B) Same metrics for human-made scenes. (C–D) Relationship between melanopsin and luminance contrast across receptive fields in natural (C) and human-made (D) environments. Error bars represent the interquartile range. In panels C and D, contrast values higher than one are not shown for graph readability purposes.

### Independence

To evaluate the dependence of melanopsin contrast with absolute excitation (intensity), we have analyzed the correlation between these two variables. We found that there is a slight but significant correlation for natural environments (*r* = −0.11, *p* < 0.05), and a similar but non-significant correlation was found for human-made environments (*r* = −0.11, *p* = 0.14) between melanopsin variables (Fig. 4). When analyzing luminance independence, we found a non-significant weak correlation for both natural (*r* = −0.01, *p* = 0.83) and human-scenes (*r* = −0.08, *p* = 0.33). These correlation values for luminance are consistent with the literature^8^. Therefore, melanopsin contrast is independent from melanopsin excitation in human-made environments. Surprisingly, melanopsin contrasts are invariable across the range of melanopsin excitation for natural scenes.

**Figure 4.**
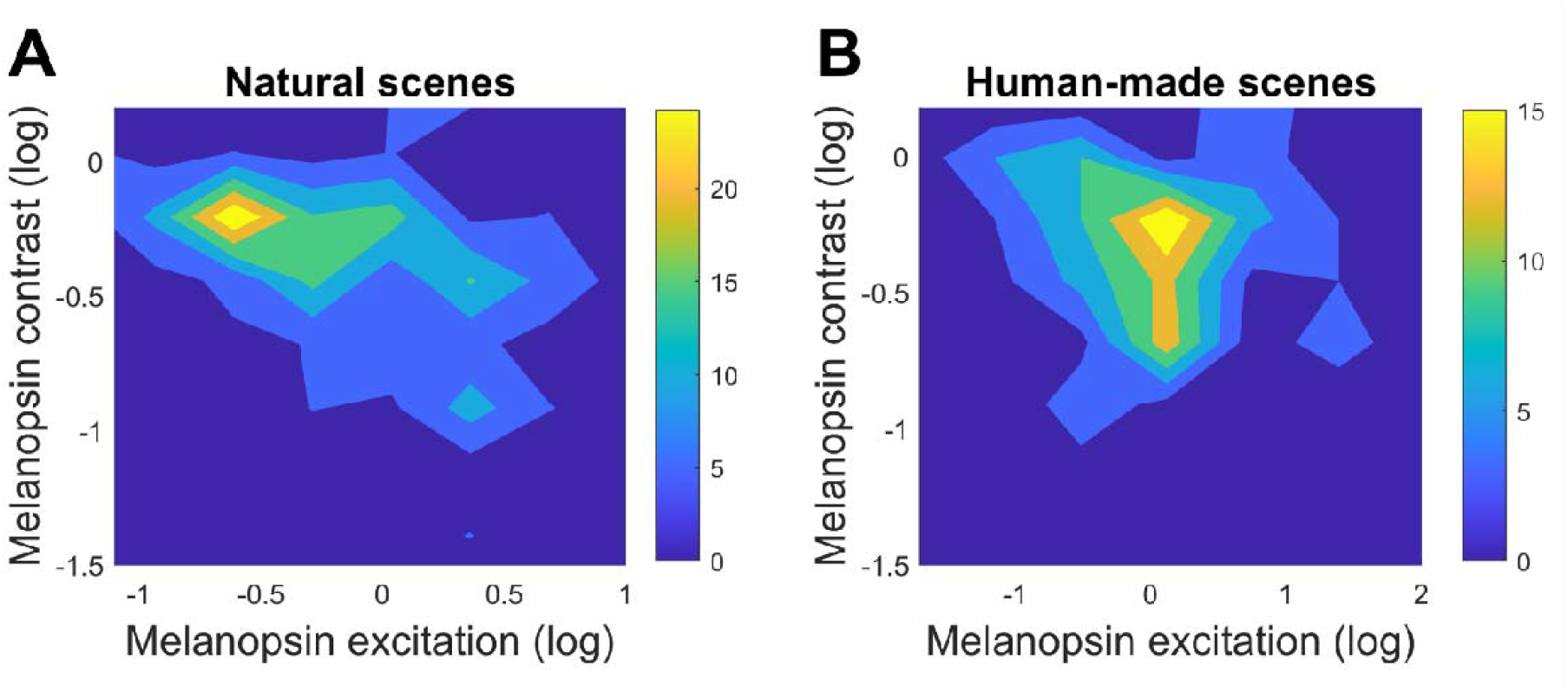
Relationship between melanopsin excitation and contrast. Colormaps showing melanopsin contrast as a function of melanopsin excitation across receptive fields in (A) natural and (B) human-made environments. Melanopsin contrast is largely independent of excitation, with only a weak correlation observed in natural scenes (r = −0.11, p < 0.05). No significant correlation is found in human-made environments.

### Effect of the size of the receptive field

For the above analyses we have used an ipRGC receptive field size of 1.37° (diameter), which corresponds to a dendritic field in the parafoveal region where ipRGCs are more abundant^35,39^. This field size is also consistent with the receptive field size used in psychophysical measurements to obtain the melanopsin contrast sensitivity function^28^. Since the size of the dendritic field of human ipRGCs increases with the eccentricity^35^, we wondered how a different receptive field size would affect the results. Therefore, we computed melanopsin excitation and contrast for natural and human-made environments considering a receptive field size of 2.4° in diameter, which corresponds to the size of ipRGC dendritic fields at the periphery (∼31° of eccentricity). We found that the distributions of the results for both sizes align quite well for excitation in both environments (Fig. 5, top panels), however a pair-wise t-test showed that the field size of 2.4° excitation was slightly but significantly higher than the data with field size of 1.37° (natural: *t* = −4.1, *p* < 0.0001; human-made: *t* = −2.08, *p* < 0.05). Regarding contrast values, the joint distribution was more disperse for both environments than excitation values (Fig. 5, bottom panels). For the natural environment, the melanopsin contrasts for the bigger field (2.4°) are significantly lower than the contrasts for the small field (*t* = 5.84, *p* < 0.0001), however this difference was not found for the human-made environment (*t* = 1.93, *p* = 0.056).

**Figure 5.**
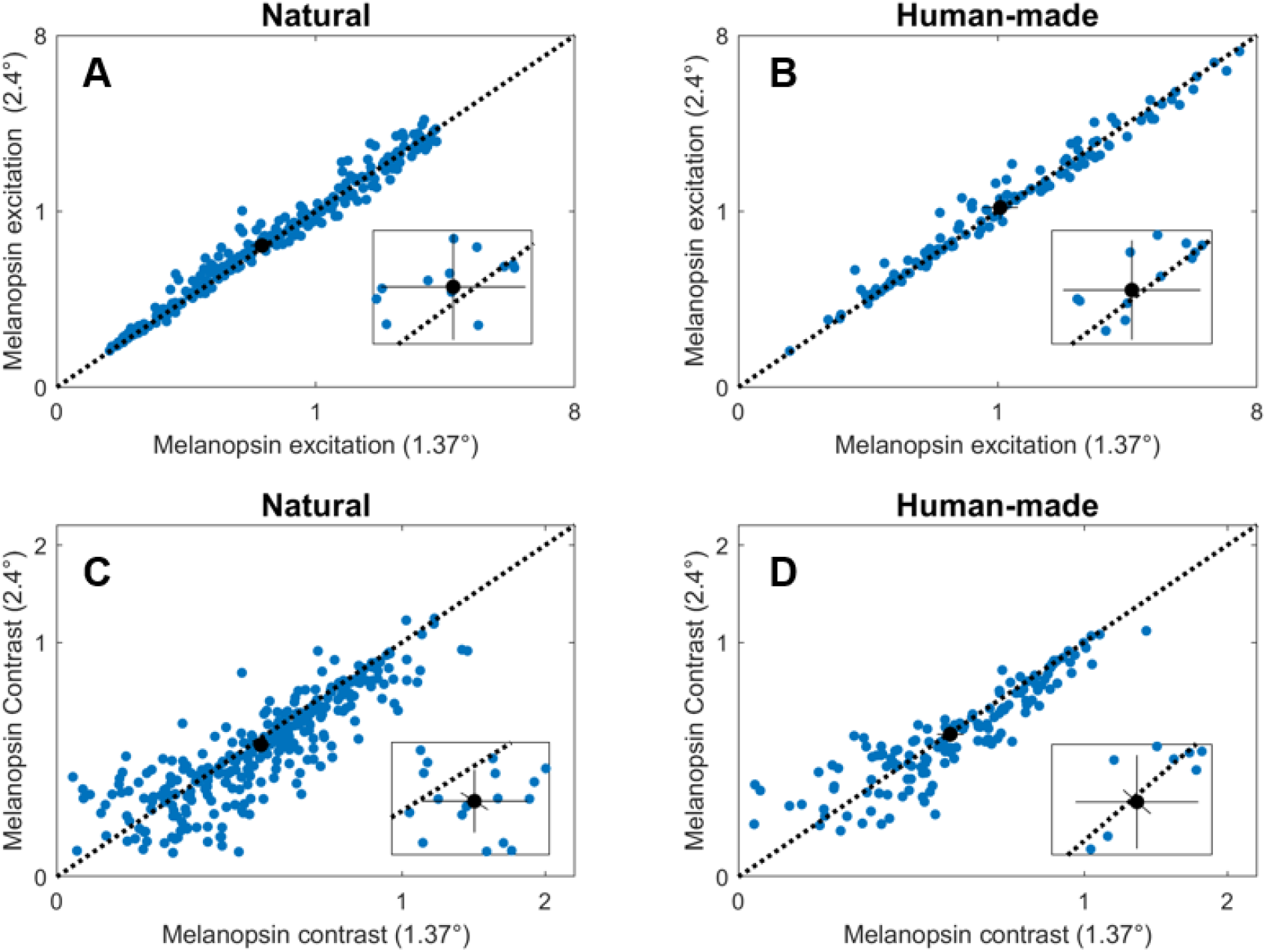
The effect of the field size in melanopsin excitation and contrast. Blue dots represent the results for each field in each image considering field diameter of 2.4° versus 1.37° for excitation in natural environments (A), excitation in human-made environments (B), contrast in natural environments (C), and contrast in human-made environments (D). The black circles represent the mean value, the black solid lines represent 95% confidence intervals^49^. The insets represent magnification of the mean value area.

### IpRGC codification

The evidence about how the ipRGCs combine intrinsic and extrinsic responses came mostly from pupillary studies that have analyzed this relationship^17^. We have used the weights obtained from a previous pupil work that isolated these responses^41^. From that study’s findings, we have set ipRGC codification following the Equation 1. In this section we called this codification as ipRGC 1. To analyze how a different weighting might affect our results we have tested a second set of weights (ipRGC 2). In this case all the weights are set to 1 (Eq. 2). We have used this second weighting selection as an approach that relies only in the normalization proposed by the CIE commission ^50^. The results of this comparison are plotted in figure 6. We found that the second codification (ipRGC 2) produced higher excitation values than ipRGC 1 for both environments (Fig. 6A), which can be evidenced by looking that no confidence interval overlapped the 45° line (Fig. 6A inset). This result was expected since overall weights are higher for ipRGC 2 than for ipRGC 1. For the ipRGC 2 codification, the human-made environments generated significantly higher excitation values than natural environments (Fig. 6B, *t* = −3.98, *p* < 0.0001) in agreement with ipRGC 1 codification. Regarding contrasts, pair-wise t-tests showed that contrasts for ipRGC 2 codification was slight but significantly higher than contrasts for ipRGC 1 in natural environment (*t* = −2.53, *p* < 0.05), but this difference was not found for human-made environment (*t* = 0.71, *p* = 0.48). This difference between ipRGC 1 and ipRGC 2 for natural environment suggest an interplay between the type of environment and the ipRGC weights of intrinsic and extrinsic inputs.

**Figure 6.**
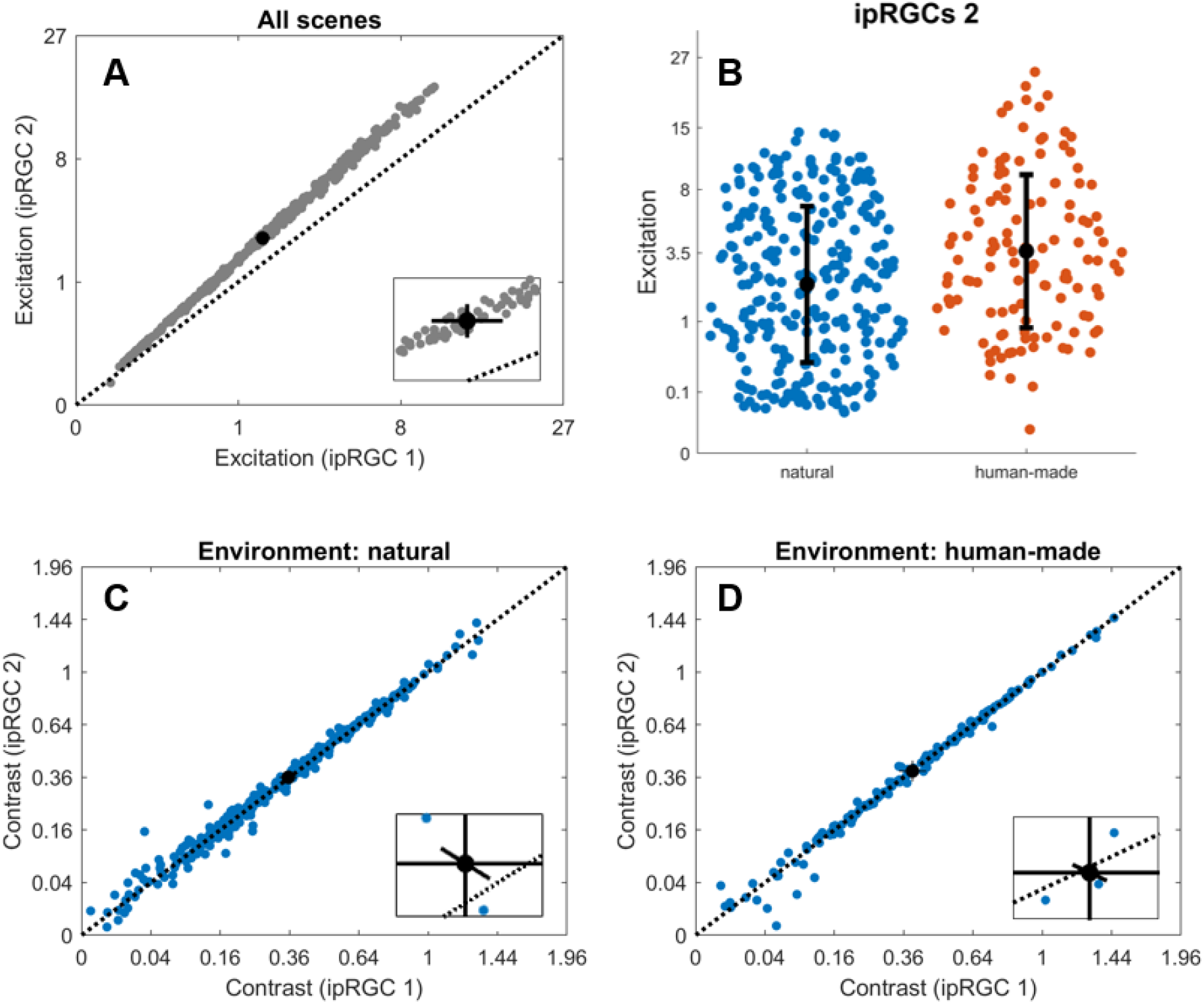
Comparison of ipRGC codification models across environments. (A) Excitation values generated by ipRGC 1 and ipRGC 2 codification models for all scenes. ipRGC 2 consistently produces higher values due to its equal weighting of photoreceptor inputs. Inset shows the deviation from the unity line. (B) Excitation values for natural and human-made scenes using ipRGC 2 model, confirming that human-made environments produce significantly higher excitation (p < 0.0001). The black circles represent the mean value and the black solid lines represent standard deviation. (C–D) Comparison of contrast values between ipRGC 1 and ipRGC 2 for natural (C) and human-made (D) environments. A significant increase in contrast is observed for ipRGC 2 in natural scenes only (p < 0.05). In A, C, and D, the black circles represent the mean value and the black solid lines represent 95% confidence intervals^49^

## Discussion

In this study, we characterized the melanopsin-mediated statistics of natural and human-made environments using hyperspectral images. Our results reveal that both the excitation and contrast properties of these environments significantly differ, with potential implications for non-image-forming processes mediated by intrinsically photosensitive retinal ganglion cells (ipRGCs).

### Human-made environments exhibit higher absolute intensities

We observed that human-made environments produce significantly higher absolute values of melanopsin excitation, ipRGC signals, and luminance when compared to natural environments. These higher levels likely result from the presence of high reflectance, geometrically regular surfaces common of built environments (like white walls or glossy surfaces), which enhance light collection and scattering. Given the central role of ipRGCs in physiological processes such as circadian entrainment, pupil regulation, mood modulation, and, even, vision, these differences in environmental light intensity may contribute to the physiological impact of urban lighting^51–53^.

### Contrast regularities differ between photoreceptor types and environments

Contrast statistics revealed environment- and photoreceptor-specific patterns. Only for natural scenes, luminance contrast exceeded both melanopsin and ipRGC contrasts when we consider the global population. However, this difference is driven by the middle and high contrast ranges. In the lower contrast range, this difference disappeared. This finding is in disagreement with the only one previous work reporting contrast regularities of melanopsin in natural scenes. In their study Allen and colleagues showed that melanopsin contrast and luminance contrast strongly correlate for the entire contrast range^54^. However, they use the same field size to compute melanopsin and luminance. If our computation uses the same filed sizes, we also obtained that strong correlation (Fig. S1). Since ipRGCs can process only coarse spatial information in comparison to the detailed information provided by luminance cells^28^, it is more accurate to include physiological-inspired field sizes to compute melanopsin and luminance contrasts.

In contrast, human-made scenes showed no significant differences between luminance and melanopsin contrasts. Since it was largely hypothesized that the retinal receptive fields are optimized to process natural stimuli^55^, this suggests that the statistical properties of artificial environments may deviate from the evolutionary constraints that shaped ipRGC function.

Previous studies have shown that natural and human-made environments differ in their first and second order luminance statistics. Natural scenes tend to exhibit broader, scale-invariant contrast distributions and greater variability in local contrast, while human-made scenes are characterized by more uniform and repetitive structures with reduced contrast variability^8,9^. These differences suggest that the early visual system, including parasol ganglion cells and potentially ipRGCs, may be adapted to efficiently encode the complex statistics of natural environments^8,56^.

### Melanopsin contrast is largely independent of excitation in nature

Our results show that melanopsin contrast is largely invariant across the range of melanopsin excitations in natural environments, with only a weak (but significant) correlation observed. This finding supports the notion that melanopsin contrast operates as a relatively independent signal dimension, enabling stable contrast detection across varied lighting conditions. Since it is thought that stable and long-term responses of ipRGCs provide unique information to visual perception complementary to the transient and adaptive rod and cone responses ^27,28,31^, such independence maybe a functional advantage for sustained physiological responses, such as circadian regulation and sustained pupil constriction. For human-made scenes, the observed relationship could be due to random chance, suggesting that natural coding feature of the melanopsin system might be affected in the build environment.

### Receptive field size influences contrast measurements

We further examined the impact of receptive field size on melanopsin contrast. While excitation values increased slightly with larger field sizes (2.4° vs. 1.37° diameter), contrast values were significantly reduced in natural environments. This reduction agrees with previous evidence of increased spatial pooling over larger retinal areas, which flatten local variations in excitation^57^. These results highlight the importance of considering spatial scale when modeling ipRGC responses, particularly given the eccentricity-dependent dendritic field sizes of these cells.

### Different ipRGC signal integration

Finally, we explored how different ipRGC codification schemes influence excitation and contrast estimates. Codification using equal weighting of cone and melanopsin inputs (ipRGC 2) resulted in higher contrasts than the biologically derived weighting (ipRGC 1) in natural environments, but not in human-made scenes. This environment-specific effect highlights the importance to confirm the weightings that ipRGCs use to combine extrinsic and intrinsic inputs.

### Limitations

Our analysis was based on a set of hyperspectral images obtained in Portugal. Although it is expected that the natural regularities that we found are kept for other environments, to confirm generality it would be necessary to analyze other environments. For this reason, we have made the scripts that we used for data analysis completely available. Also, some of the analyzed scenes contained both built and natural structures, the decision to tag them as natural or human-made was made arbitrarily considering the predominant surfaces in those scenes. Finally, other models were also proposed to take into account the combination of intrinsic and extrinsic inputs in ipRGCs (for example^20^). Testing of further models was outside the scope of this study.

## Conclusion

This study provides the first systematic analysis of melanopsin and ipRGC signal statistics across natural and human-made environments using hyperspectral imaging. We found that human-made environments exhibit significantly higher absolute intensities across melanopsin, ipRGC, and luminance signals, while natural environments preserve a distinct contrast structure, with luminance contrasts dominating in higher ranges over melanopsin contrasts. Melanopsin contrast was largely independent of absolute excitation, indicating a robust encoding mechanism across lighting conditions. Additionally, receptive field size significantly influenced contrast estimates, and variations in ipRGC codification models revealed an interaction between signal weighting and environmental context. Together, these findings highlight the unique statistical features of natural environments to which the melanopsin system may be evolutionarily tuned and suggest that deviations in modern visual environments could alter ipRGC-mediated physiological functions.

## Methods

### Hyperspectral Image Dataset

We used a publicly available dataset of hyperspectral images of natural and human-made environments under daylight illumination, previously published by Foster and Nascimento^46,47^. A total of 31 images were included in the analysis: 21 scenes characterized as natural (e.g., vegetation, natural terrain) and 10 scenes as human-made (e.g., urban structures). All images were acquired in outdoor settings in Portugal and cover the visible spectrum from 400 to 700 nm.

### Spectral Preprocessing and Excitation Computation

Each hyperspectral image was interpolated to a uniform wavelength sampling interval of 5 nm to ensure consistent spectral resolution across the dataset. For each pixel, the radiance spectra were convolved with the CIE physiological cone fundamentals^58^ and melanopsin spectral sensitivity^50^ (Fig. 1A, inset) to compute spatially resolved maps of photoreceptor excitation: L-, M-, and S-cones, and melanopsin. The melanopsin photopigment has a peak of absorbance around 480 nm^14^, however the spectral sensitivity peak at corneal level shifts due to intraocular media filtering^50^. In comparison, human cones are classified in three subtypes, L, M and S-cones, with peak wavelength around 565 nm, 535 nm, and 419 nm, respectively^44^. Photopic luminance was computed as the weighted sum of L and M cone excitations using the CIE V(λ) function (Fig. 1A, inset). Excitations were expressed in α-opic equivalent luminance units.

### ipRGC Signal Codification

To estimate ipRGC responses, we modeled two types of codification based on known physiological inputs. The first model (ipRGC 1) used a weighted sum of photoreceptor inputs derived from human pupillometry studies^41^ (Equation 1):

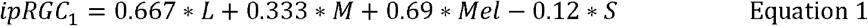

Where ipRGC_1_ is the possible response of combined extrinsic (L, M and S cones) and intrinsic (Mel, melanopsin) inputs, following the first model.

A second model (ipRGC 2) used equal weighting for L, M, and melanopsin inputs and subtractive weighting for S-cones (Equation 2):

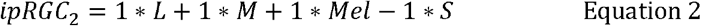

Where *ipRGC*_2_ is the possible response of combined extrinsic (L, M and S cones) and intrinsic (Mel, melanopsin) inputs, following the second model.

These models were applied pixel-wise to generate spatial maps of ipRGC activation.

### Receptive Field Modeling and Contrast Calculation

Receptive fields were simulated using circular, symmetric, raised cosine window functions (Equation 3)^8,56^, centered on non-overlapping patches within each image. Two receptive field diameters were modeled to reflect ipRGC dendritic field sizes in the parafovea (1.37°) and periphery (2.4°)^35^. For luminance calculations, a smaller receptive field of 0.36° diameter was used, consistent with parasol ganglion cell physiology in the parafoveal region^59^. To compute sizes we used a retinal magnification factor of 0.291 mm/deg^28,60^.

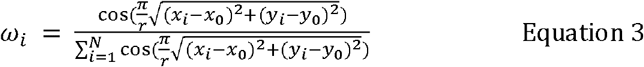

Where ω_i_ the weight from the windows function, r is the radius of the patch, (*x*_i_, *y*_i_) is the location of the i^th^ pixel in the patch, and (*x*_0_, *y*_0_) is the location of the center of the patch.

**Local excitation** was defined as the weighted mean response within a receptive field (Equation 4). In addition, **between-patches contrast** was calculated as the absolute Michelson contrast of the local excitation values across all patches within an image (Equation 5).

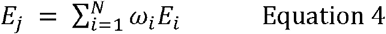

Where *E*_*j*_ is the local excitation of the j^th^ patch, and *E*_*i*_ is the excitation at the i^th^ pixel in the patch.

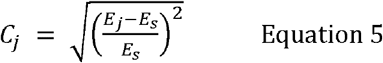

Where *C*_*j*_ is the between-patches contrast of the j^th^ patch, and *E*_*s*_ is the mean excitation of the patches in the s^th^ scene.

## Statistical Analysis

All contrast and excitation values were aggregated across scenes within each environment type. Group comparisons (natural vs. human-made) were assessed using analysis of variance, paired and independent-samples t-tests, or two-sample F test for equal variances, depending on the analysis required. Pearson correlation was used to evaluate the relationship between excitation and contrast measures. Analysis and plotting of mixed confidence intervals in scatter plots were done with the software provided by Dr. Alexander Schütz^61^. Statistical significance was set at p < 0.05, and all analyses were conducted using MATLAB (MathWorks) and GraphPad Prism (GraphPad Software). The scripts used in this work are publicly available in https://github.com/francisco-diaz-barrancas/HyperMelanopsin.

## Code Validation

To validate the contrast computation pipeline, we generated artificial scenes using silent substitution techniques^62,63^, containing either melanopsin-only or luminance-only contrast stimuli. These stimuli confirmed that our code correctly isolated and quantified the intended photoreceptor-specific contrasts without spurious activation of other channels (Fig. S2).

## Acknowledgements

This study has received funding from the German Research Foundation (DFG) [222641018 - SFB/TRR 135 TPs C2 and B2], the European Research Council (ERC) under the European Union’s Horizon 2020 research and innovation programme [project “SENCES” number: 101001250, and project “COLOR3.0” number 884116], the Agencia I+D+i [PICT2019-03673], the Consejo Nacional de Investigaciones Científicas y Técnicas [PIP-2721], the Regional Ministry of Education, Science and FP of the Government of Extremadura [GR24030], and the “European Union NextGenerationEU/PRTR” [C110.23].

