## Supplementary material for "Melanopsin-mediated signals in natural and human-made environments"

**Table S1. Name and sources of the hyperspectral images used in this study.**

| Environment | Name | Associated article | obtained from: |
|---|---|---|---|
| Natural | Bom_Jesus_Bush | Nascimento, Amano & Foster (2016)[1] | A |
| Natural | Bom_Jesus_Marigold | Nascimento, Amano & Foster (2016)[1] | A |
| Natural | Bom_Jesus_Red_flower | Nascimento, Amano & Foster (2016)[1] | A |
| Natural | Bom_Jesus_Ruin | Nascimento, Amano & Foster (2016)[1] | A |
| Natural | Gualtar_Orange_Trees | Nascimento, Amano & Foster (2016)[1] | A |
| Natural | Lillies_Closeup | Nascimento, Amano & Foster (2016)[1] | A |
| Natural | Lilly_Closeup | Nascimento, Amano & Foster (2016)[1] | A |
| Natural | Ruivaes_Fern | Nascimento, Amano & Foster (2016)[1] | A |
| Natural | Ruivaes_Ruin | Nascimento, Amano & Foster (2016)[1] | A |
| Natural | Sameiro_Bark | Nascimento, Amano & Foster (2016)[1] | A |
| Natural | Sameiro_Branch | Nascimento, Amano & Foster (2016)[1] | A |
| Natural | Sameiro_Forest | Nascimento, Amano & Foster (2016)[1] | A |
| Natural | Sameiro_Glade | Nascimento, Amano & Foster (2016)[1] | A |
| Natural | Sameiro_Leaves | Nascimento, Amano & Foster (2016)[1] | A |
| Natural | Sameiro_Trees | Nascimento, Amano & Foster (2016)[1] | A |
| Natural | Sete_Fontes_Rock | Nascimento, Amano & Foster (2016)[1] | A |
| Natural | Tibaes_Garden | Nascimento, Amano & Foster (2016)[1] | A |
| Natural | Yellow_Rose | Nascimento, Amano & Foster (2016)[1] | A |
| Natural | levada_1411 | Foster, Amano & Nascimento (2016)[2] | B |
| Natural | nogueiro_1441 | Foster, Amano & Nascimento (2016)[2] | B |
| Natural | sete_fontes_1438 | Foster, Amano & Nascimento (2016)[2] | B |
| Human-made | Braga_Grafitti | Nascimento, Amano & Foster (2016)[1] | A |
| Human-made | Gualtar_Columns | Nascimento, Amano & Foster (2016)[1] | A |
| Human-made | Gualtar_Villa | Nascimento, Amano & Foster (2016)[1] | A |
| Human-made | Souto_Farm_Barn | Nascimento, Amano & Foster (2016)[1] | A |
| Human-made | Souto_Roof_Tiles | Nascimento, Amano & Foster (2016)[1] | A |
| Human-made | Souto_Wood_Pile | Nascimento, Amano & Foster (2016)[1] | A |
| Human-made | Tenoes_Wall | Nascimento, Amano & Foster (2016)[1] | A |
| Human-made | Tenoes_Wall_Closeup | Nascimento, Amano & Foster (2016)[1] | A |
| Human-made | Tibaes_Corridor | Nascimento, Amano & Foster (2016)[1] | A |
| Human-made | gualtar_Steps | Nascimento, Amano & Foster (2016)[1] | A |

A) https://sites.google.com/view/sergionascimento/home/scientific-data/hyperspectral-images-for-spatial-distribution-of-local-illumination-2015. B) https://sites.google.com/view/sergionascimento/home/scientific-data/time-lapse-hyperspectral-radiance-images-2015.

**Table S2. Summary of the statistics values from t-tests of natural versus human-made environments.**

| | Excitation | | | | Contrast | | | |
|---|---|---|---|---|---|---|---|---|
| | mean | | variance | | mean | | variance | |
| | t | p | f | p | t | p | f | p |
| Melanopsin | -5.390 | 1.17E-07 | 0.890 | 0.439 | -1.095 | 0.274 | 1.000 | 0.999 |
| ipRGC$_1$ | -4.299 | 2.14E-05 | 1.015 | 0.938 | -1.235 | 0.217 | 0.931 | 0.592 |
| Luminance | -3.667 | 2.78E-04 | 0.977 | 0.861 | -0.615 | 0.539 | 0.929 | 0.583 |

**Figure S1. Melanopsin versus luminance contrast for the same field size.** These scatter plots show the high correlation between melanopsin and luminance contrasts in both environments when the same, but not real, receptive field size is used (1.37°). These results showed the importance of using the proper receptive field sizes when comparing luminance and melanopsin signals.

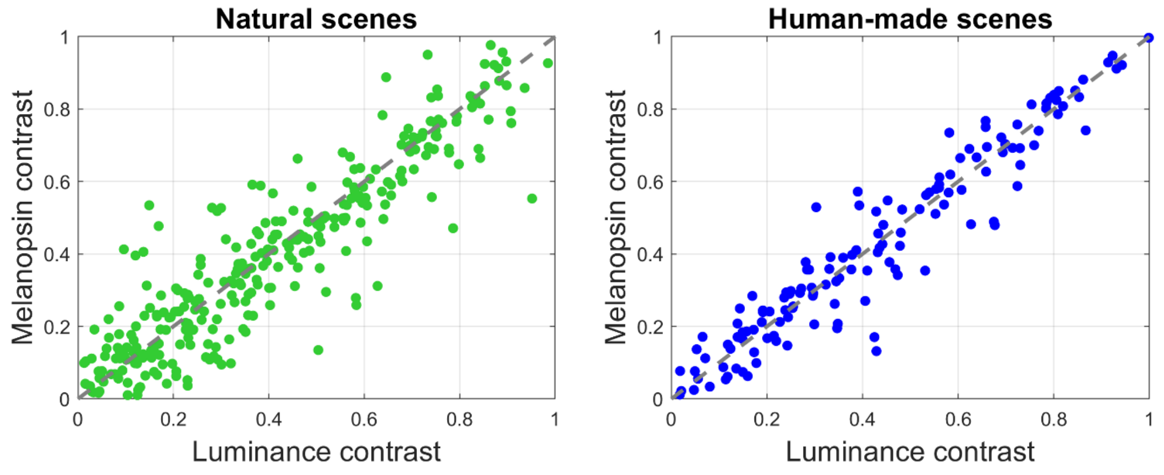


**Figure S2. Computation validation.** To validate the computation, we generated artificial hyperspectral scenes. Three of these scenes were generated using the silent substitution method[3,4], which allows to generate stimulation that selectively stimulate individual or combined photoreceptor excitations. These scenes were divided in sixteen patches, each patch could contain a positive or negative contrast for an intended excitation (luminance or melanopsin). The first scene only differed in luminance (combined L and M-cone excitation change, while maintaining stable excitation of melanopsin, S-cones and rods). The second scene only differed in melanopsin excitation (while maintaining stable excitation of rods, L-, M- and S-cones). The third scene completely produced an equal energy spectrum (no contrast changes for any photoreceptor). The fourth scene replicated the output values of a five-primary photostimulator, which is a device built to selectively stimulate melanopsin in laboratory settings[5]. We computed contrast for these four artificial scenes with the same codes that we used for the main analyses. As expected, our contrast computation generated changes only for the intended excitation. These results validated our computations and ensured non-spurious signals.

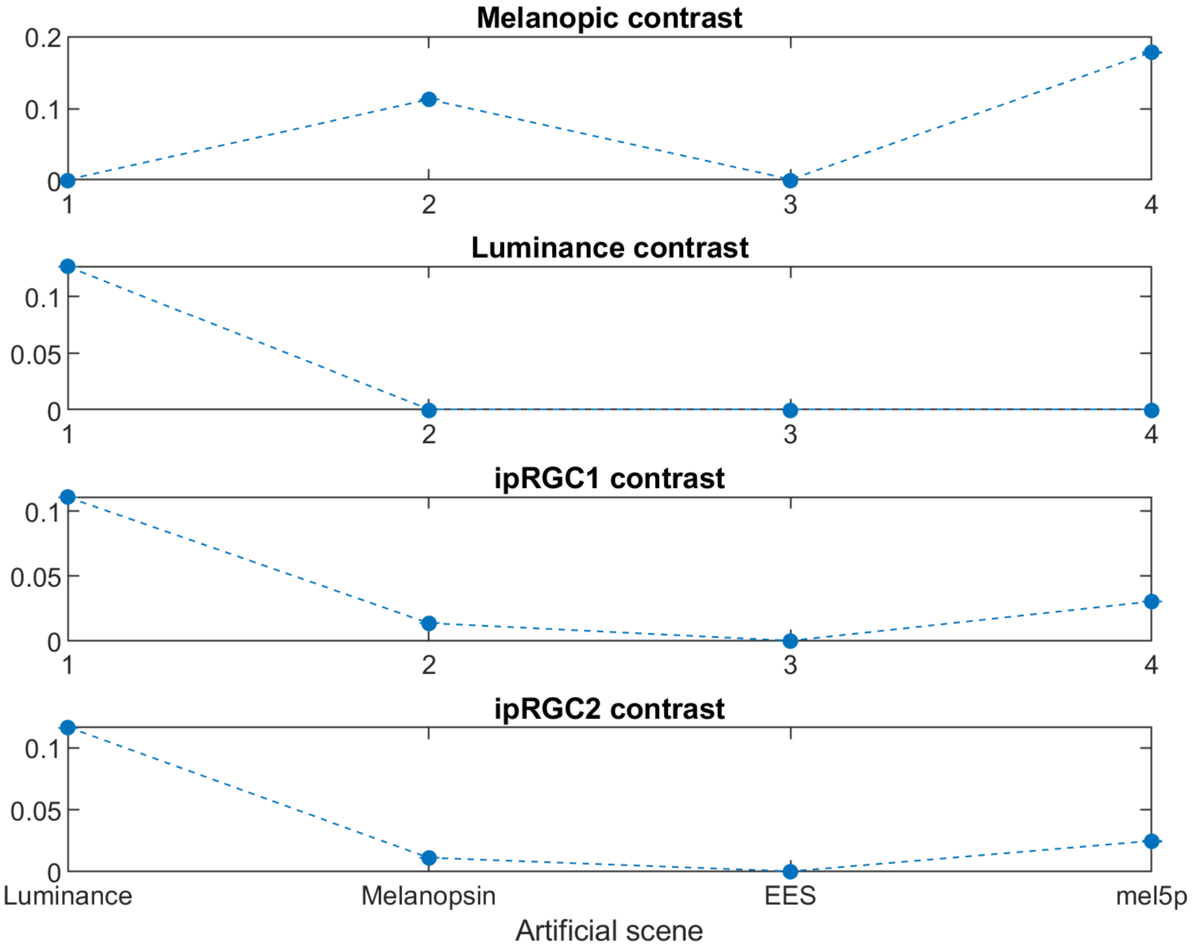
